# The *Salmonella* effector SspH2 disrupts F-actin cytoskeletal organization to inhibit macrophage migration

**DOI:** 10.64898/2026.10.14.759247

**Authors:** Una Deng, Melissa Gonzalez, Rose Mera, Ji’Shuri Johnson, Alondra Mancillas, Natasha Sanchez, Jayme Angelo, Leah Hernandez, Ramiro Hernandez, Kristen Weber, Manuel Amieva, Denise M Monack, Laura M McLaughlin

## Abstract

The facultative intracellular pathogen *Salmonella enterica* serovar Typhimurium (*S. Typhimurium*) causes gastroenteritis in humans and persistent infections in animals, enabling widespread environmental reservoirs. Intracellular survival of *S. Typhimurium* depends on a suite of type III secretion system effectors encoded within *Salmonella* Pathogenicity Island 2 (SPI-2), many of which also impair host cell migration. Among these, SspH2 is a novel E3 ubiquitin ligase (NEL) effector, but how it inhibits host cell movement has remained unclear. By using a combination of time-lapse microscopy and wound-healing assays, we show that SspH2 is required for *S. Typhimurium* to fully suppress migration of macrophage-like RAW264.7 cells and that conditional expression of SspH2 alone is sufficient to restrict their movement. SspH2 expression led to a marked reduction in F-actin and fewer cells containing higher-order F-actin structures, indicating cytoskeletal remodeling. In contrast, the catalytically inactive mutant SspH2C580A had no effect, demonstrating that E3 ligase activity is essential for these phenotypes. Knockdown of the host cytoskeletal factor vinculin, a core component of podosomes and focal adhesions, mimicked SspH2’s effect on cell migration. When vinculin expression was reduced, SspH2 did not further inhibit migration, suggesting that SspH2 requires vinculin-associated F-actin-rich structures to block cell migration. These findings reveal a new pathway by which *S. Typhimurium* limits host cell movement, potentially enhancing immune evasion and fitness within the host.

## Introduction

Non-typhoidal *Salmonella enterica* serovars are a major cause of bacterial foodborne gastroenteritis, responsible for an estimated 93.8 million cases and 155,000 deaths globally each year [1, 2]. One of the most common serovars in the United States, *Salmonella enterica* serovar Typhimurium (*S. Typhimurium*), is a facultative intracellular pathogen that causes gastroenteritis in humans and can systemically infect and persist in animals such as mice [3]. A feature of *S. Typhimurium* pathogenesis is its deployment of two type III secretion systems (T3SSs), encoded on *Salmonella* pathogenicity islands 1 and 2 (SPI-1 and SPI-2). The SPI-1 T3SS primarily facilitates bacterial invasion into non-phagocytic cells [4], while the SPI-2 T3SS translocates effector proteins into host cells following invasion or phagocytosis. These effectors remodel the host cell to form a *Salmonella*-containing vacuole (SCV) that supports bacterial replication [5] and can also limit antigen presentation by phagocytes, delaying the host immune response [6–8].

One strategy used by *S. Typhimurium* to subvert host immunity is to interfere with the regulation of host cell migration [9–14]. The SPI-2 effector SseI (SrfH) restricts macrophage and dendritic cell chemotaxis, allowing the pathogen to evade immune clearance and persist in its murine host [9]. A screen for SPI-2 effector mutants revealed several additional effectors that modulate host immune cell migration, including the novel E3 ubiquitin ligase (NEL) family protein, SspH2 [10]. Recent work demonstrated that SspH2 inhibits cell migration via its E3 ligase activity when ectopically expressed in human embryonic kidney (293T) cells [14], where it binds and poly-ubiquitinates the host factor LMO4, a transcriptional co-activator that can also localize to the leading edge of migrating cells [15].

SspH2 was previously shown to contribute to the virulence of *S. enterica* serovars Typhimurium [16] and Enteritidis [17] in vivo, although the underlying mechanism remains unclear. Structurally, SspH2 contains an N-terminal leucine-rich repeat (LRR) domain that mediates substrate binding and a C-terminal NEL domain harboring a conserved catalytic cysteine (C580) essential for E3 ligase activity [18]. Once inside host cells, SspH2 is S-palmyloylated, which anchors it to the cytoplasmic membrane [19]. SspH2 also binds to cytoskeletal proteins such as α-filamin and profilin-1, interactions that, although not linked to ubiquitination, suggest a role in cytoskeletal regulation [20–22]. In addition, SspH2 can ubiquitinate the host innate immune receptor NOD1 [23, 24], and inhibits antigen presentation when expressed in professional antigen-presenting cells [6]. Whether SspH2 manipulates immune cell migration as part of its immune evasion strategy remains an open question.

Migration of macrophages and dendritic cells is essential for host defense, as these cells must traffic to and from sites of infection. In Wiscott-Aldrich Syndrome, a genetic immunodeficiency, macrophages and dendritic cells exhibit defective migration [25, 26] and lack podosomes [27, 28]–F-actin-rich structures critical for adhesion and migration [29]. Vinculin, a shared component of podosomes and focal adhesions, is a mechanosensitive factor that forms a complex with other adhesion proteins to link F-actin to the extracellular matrix via integrin binding [30–33]. Pathogens such as *Shigella* [34–36] and *Chlamydia* [37] exploit vinculin-associated pathways to alter host cell adhesion and movement, thereby promoting intracellular persistence. Whether *Salmonella* SPI-2 effectors similarly target vinculin-dependent structures has not been established.

Here, we show that the SPI-2 effector SspH2 inhibits macrophage migration by disrupting actin cytoskeletal organization. Using murine RAW264.7 macrophage-like cells, we demonstrate that SspH2 is sufficient to reduce migration speed without significantly impairing chemotaxis toward CCL5 and that its expression decreases both total F-actin and the number of cells with higher-order F-actin-rich structures arranged into rosette/ring-like formations. These effects require SspH2’s E3 ubiquitin ligase activity, indicating that *S. Typhimurium* co-opts ubiquitin-dependent pathways to perturb F-actin dynamics and restrict host cell motility. Finally, we found that SspH2-mediated inhibition of migration required the presence of the podosome-associated protein vinculin, suggesting that SspH2 may interfere with these F-actin-rich structures to modulate cell migration.

## Materials and methods

### Mammalian cell culture and *S. Typhimurium* infection

The RAW264.7 (ATCC TIB-71) cell line (and all cell lines derived from this line) was cultured in RPMI1640 supplemented with 10% fetal bovine serum (FBS) at 37°C with 5% CO_2_. The RAW264.7 caspase 1/11-/-cell line was obtained from Denise Monack (Stanford) and was described previously [38]. *S. Typhimurium* SL1344 and all SPI-2 effector mutants derived from this strain mentioned here were obtained from Denise Monack and described previously [10]. RAW264.7 caspase 1/11-/-cells were infected with *S. Typhimurium* that was grown overnight in Luria Broth (LB) at 37°C without agitation to induce SPI-1 and increase infection efficiency [4], whereas wild-type RAW264.7 cells were infected with *S. Typhimurium* that was grown overnight with agitation. Bacterial cultures were added to monolayers of RAW264.7 caspase 1/11-/-cells at a multiplicity of infection (MOI) of 30:1, and extracellular bacteria were killed 30 minutes later by replacing the medium with that containing 100 µg/mL gentamincin. After 1.5 h, the gentamicin concentration was reduced to 10 µg/mL.

### Construction of SPI-2 effector-expressing RAW264.7 cell lines

The RAW264.7 cell line stably expressing the reverse tetracycline repressor-VP16 transactivator fusion protein (pLenti CMV rtTA3 Blast (w756-1), Addgene, #26429) was generated using lentiviral delivery. Stable transductants were selected with blasticidin and dilution plating was used to isolate a clonal cell line. This reverse tetracycline-controlled transactivator (rtTA) [39] expressing RAW264.7 cell line was used as the parental cell line for all other SPI-2 effector-expressing RAW264.7 cell lines described here. To create these SPI-2 effector-expressing cell lines, the *sspH2* and *sseI* genes were amplified from *S. Typhimurium* SL1344 gDNA and the *gfp* gene was amplified from the pFPV25.1 (Addgene, #20668) plasmid DNA. These genes were then inserted into the pLenti CMVTRE3G Puro Dest (w811-1) vector (Addgene, #27565, digested with EcoRV) after a doxycycline-inducible promoter using the Gibson cloning kit from NEB [40], thereby tagging the SPI-2 effector at the C-terminus with GFP. Primers were developed with the help of Benchling.com’s Gibson cloning wizard and are shown in Table 1. The resulting plasmid constructs were amplified in recombination-deficient *Escherichia coli* (NEB stable *E. coli*, NEB) and verified by DNA sequencing (Elim Biopharm).

**Table 1:**
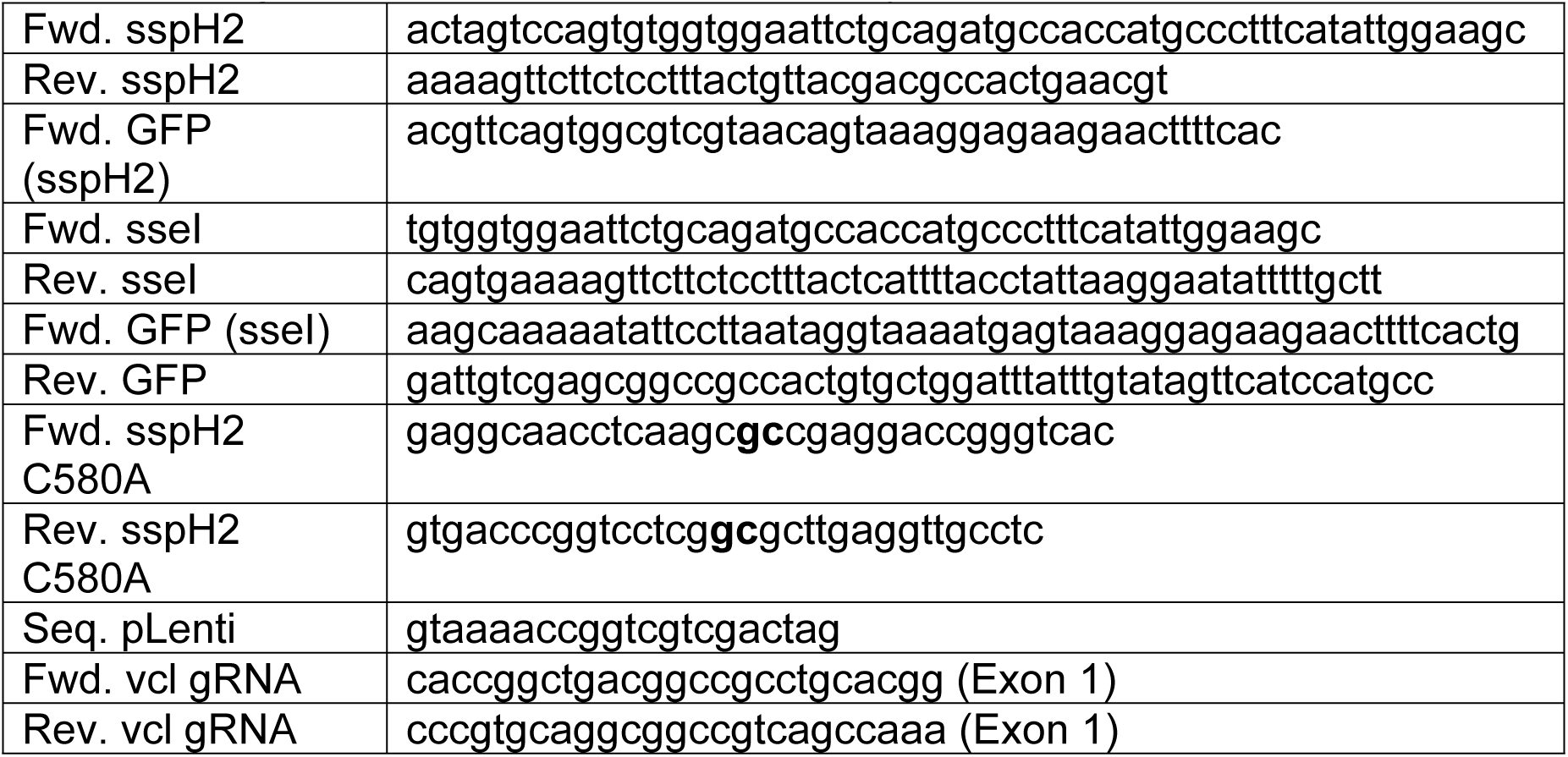
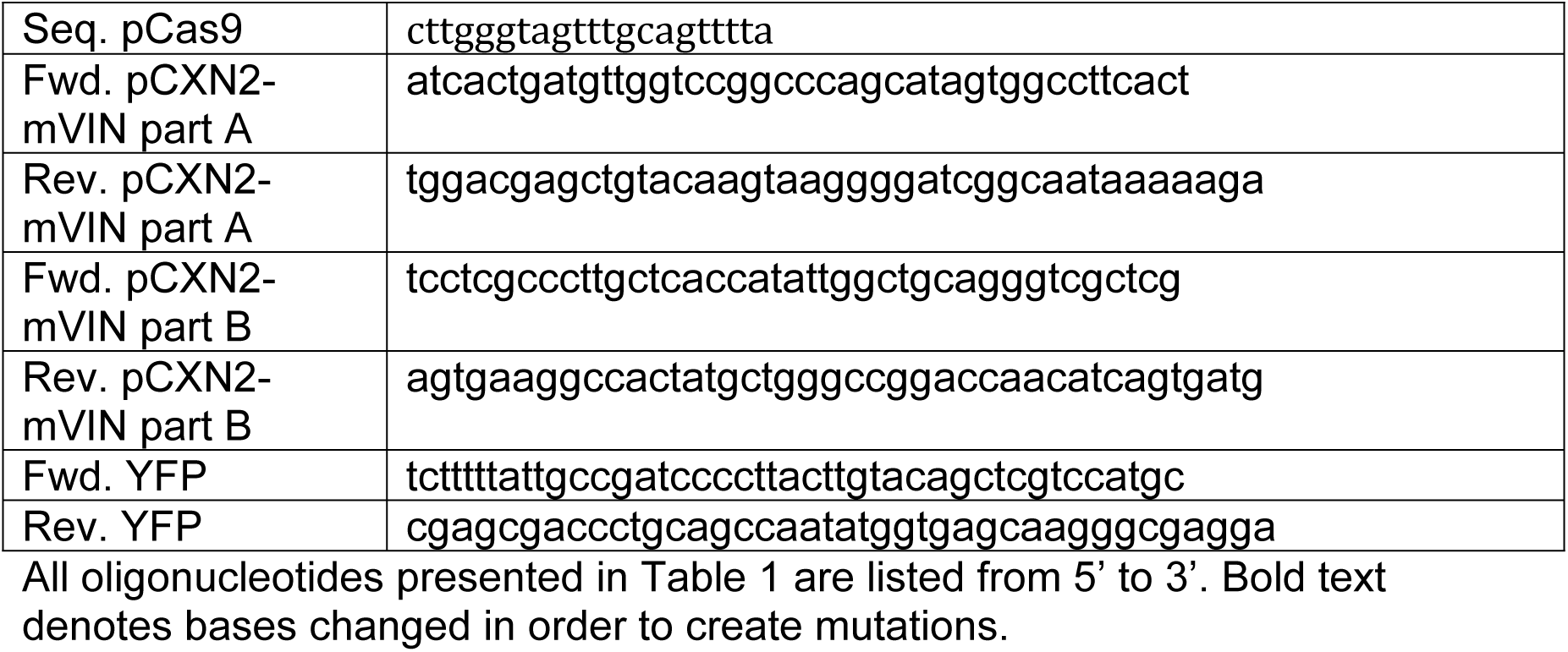
Oligonucleotides used in this study.

A pLenti vector carrying a C580A mutant form of SspH2 tagged with GFP was created by site-directed mutagenesis using the pLenti-SspH2-GFP vector made here as the template for PCR amplification. Two PCR products were generated, one using the for. sspH2 and rev. sspH2 C580A primers and another using the for. sspH2 C580A and rev. GFP primers (Table 1, bold denotes mutation). These two DNA fragments were combined with more of the pLenti CMVTRE3G Puro Dest (w811-1) vector (Addgene, #27565, digested with EcoRV) using the Gibson cloning kit from NEB.

The resulting pLenti-SPI-2 effector plasmids were then transfected along with the helper plasmids (pCMV-VSV-G (Addgene #8454), pCMV-dR8.2_dvpr (Addgene #8455) and pAdVAntage (Promega)) into 293T (ATCC #CRL-3216) lentiviral packaging cells using TransFectin reagent (BioRad). Transducing particles were harvested and used to transduce the parental RAW264.7 cell line. Stable transductants were selected and maintained in cell medium supplemented with 5 µg/mL puromycin. As a control, a RAW264.7 cell line that expresses mCherry was produced by similarly using a pLenti-mCherry vector (CM Dovey). Finally, the amount of doxycycline necessary to induce SPI-2 effector-GFP expression (as evidenced by detectable GFP fluorescence) was determined to be between 10 and 100 ng/mL for each SPI-2 effector-expressing cell line. Unless otherwise specified, 50 ng/mL doxycyline was used to induce SPI-2 effector expression in all experiments presented here.

### Cell migration

#### In vitro scratch assay

Essentially as previously described [41–43], a monolayer of the desired RAW264.7 cell line (with or without infection by *S. Typhimurium*, as described above) was scratched using a 200µl pipette tip and three points along this line were labeled on the back of the plate. After changing the medium (adding doxycycline where necessary), images of the scratch at each point were taken immediately with an inverted light microscope (Olympus) at 100X magnification and then again one or two days later. Cell migration into the scratch was determined by measuring the change in scratch width (Fiji).

#### Time-lapse microscopy

Subconfluent RAW264.7 cells (with or without doxycycline) were seeded onto coverglass (coated with 1% BSA for one hour at 37°C) and imaged at regular time intervals (as described in the text) at 600X, during which the cells were kept in a humidified chamber at 37°C with CO_2_ (Nikon). Fluorescence and DIC images were compiled into videos and cells were tracked to characterize cell movement behavior. Cells were tracked manually or automatically using the Trackmate plugin from Fiji.

#### Under-agarose assay

Essentially as previously described [43,44], 1% cell culture grade agarose in PBS was combined with cell medium and poured over coverglass (coated with 1% BSA). Two 3.5 mm diameter holes were punched out of the agarose 2 mm apart and 10 µl of 500 ng/mL CCL5 was added to one of the wells. After incubating the agarose plate at 37°C for 10min, 40,000 cells (a mixture of RAW264.7 cells expressing a given SPI-2 effector tagged with GFP and RAW264.7 cells expressing mCherry alone as an internal control) were seeded into the other well and allowed to incubate for 24 hours (in the presence of doxycycline). Images were taken of cells on the side of the well towards and away from the well containing the CCL5 chemokine at 600X magnification (Nikon), and chemotaxis was measured as the number of cells migrating under the agarose toward the chemokine minus the number of cells migrating under the agarose away from the chemokine. Chemotaxis of cells expressing SPI-2 effectors tagged with GFP was then normalized to chemotaxis of the mCherry-expressing internal control cells.

### F-actin characterization

#### Phalloidin staining

RAW264.7 cell lines were seeded onto BSA-coated coverslips. Once the cells had attached, 10 ng/mL doxycycline was added to induce SPI-2 effector expression. The following day, the cells were fixed in 2% paraformaldehyde in PBS, and then stained for F-actin using a 100-fold dilution of phalloidin-Alexa Fluor 660 for one hour at room temperature. After washing away excess stain, the coverslips were mounted in VectaShield with DAPI and subjected to confocal microscopy using 630X magnification (Zeiss). The intensity of each cell’s phalloidin staining was measured using Volocity software.

#### F-actin measurement in live cells

These SPI-2 effector-expressing cell lines were transfected with pmCherry-LifeAct-7 (Addgene #54491) and stable transfectants were selected in G418. These RAW264.7 cell lines were subjected to time-lapse microscopy as described above and F-actin-associated fluorescence was tracked and quantified using CellTK software [45]. Images of these cells were also used to quantify the number of cells that contained rosette/ring-like formations of punctate F-actin-rich structures at zero and one day after starting doxycycline treatment. Rosette/ring structures were defined as a circular arrangement of at least 10 individual F-actin puncta or as a continuous ring of F-actin in a given cell.

#### F-actin fractionation

Confluent 60 mm dishes of SspH2- or SspH2C580A-expressing cell lines were treated with or without doxycycline (or 10 µM cytochalasin D as a positive control), and one day later these cells were harvested on ice in F-actin stabilizing buffer (20 mM HEPES pH 6.8, 100 mM HaF, 50 mM KCl, 2 mM MgCl_2_, 0.5% triton X-100, 1X protease inhibitor cocktail (cOmplete, Roche) and 1 mM ATP) by scraping. After shearing the DNA, cell extracts were centrifuged at 10,000g for 5 minutes to remove nuclei and mitochondria, and then supernatants were ultra-centrifuged at 100,000g for one hour at 4°C. The supernatants (G-actin) and pellets (F-actin) were then subjected to SDS-PAGE and western blotting using monoclonal anti-β-actin antibody (Sigma, clone AC-15) and anti-mouse secondary antibody conjugated to Alexa Fluor 800. Bands corresponding to G- and F-actin were quantified using the LI-COR Odyssey imaging system.

### Vinculin gene disruption in SspH2-expressing cell lines

Using Benchling.com’s CRISPR guide, a small gRNA sequence targeting exon one of the murine vinculin gene (*vcl*) was chosen to use for CRISPR-targeting. Using the Golden Gate cloning kit from NEB, small oligonucleotides containing the gRNA sequence with the required overhangs (Table 1) were combined with the Cas9-expression vector pSpCas9(BB)-2A-Hygro (Addgene #127763), already digested with BbsI. The resulting *vcl*-gRNA/Cas9 expression construct was amplified in NEB stable *E. coli* and verified by DNA sequencing (Elim Biopharm). These plasmids were then transfected into SspH2-GFP-expressing RAW264.7 cells using TransFectin (BioRad) and stable transfectants were selected in hygromycin. Knockdown of vinculin expression was confirmed by immunoblotting. Briefly, protein was extracted from RAW264.7 cells (with or without CRISPR-targeting of *vcl*) using RIPA buffer (50mM Tris pH 7.4, 150mM NaCl, 0.1% sodium dodecyl sulfate, 0.5% sodium deoxycholate, 1% Triton X-100), separated by SDS-PAGE and blotted onto polyvinylidene fluoride (PVDF) membrane. Vinculin protein was detected using anti-vinculin antibody (Sigma, clone hVIN-1).

### Vinculin gene complementation

The *yfp* gene was inserted into the vinculin expression vector pCXN2-mVIN (Addgene #11776) so that both YFP and vinculin would be constitutively expressed in all transfected cells. Briefly, the pCXN2-mVIN plasmid sequence was amplified in two parts (“A” and “B”) and a third DNA fragment containing the *yfp* gene sequence was amplified from pcDNA3-YFP (Addgene #13033). These fragments were combined to form the p*Vcl* expression plasmid using the Gibson cloning kit from NEB. Resulting plasmids were verified by whole plasmid sequencing (Elim Biopharm). Transfectants were sorted on the basis of YFP expression (Sony SH800) and YFP+ cells were then seeded onto coverglass and subjected time-lapse microscopy as described above.

### Statistical methods

Unless otherwise stated, all statistical tests were carried out in Graphpad Prism or Microsoft Excel using a two-sample two-tailed Student’s t-test with an alpha level of 0.05. All graphs depict means of at least three independent experimental replicates (N given in figures) and all error bars represent standard errors of the mean (SEM).

## Results

### *S. Typhimurium* inhibits macrophage migration through SPI-2 effector SspH2

*S. Typhimurium* can manipulate the migration of primary macrophages and dendritic cells through several SPI-2 effectors [9, 10]. To test whether this activity extends to the murine macrophage-like cell line RAW264.7, we performed an in vitro scratch assay (Fig 1). To maximize infection efficiency, *S. Typhimurium* was grown overnight without agitation to induce SPI-1 prior to infection, thereby increasing bacterial invasiveness [4]. To prevent inflammasome-dependent cell death during infection, we used a RAW264.7 cell line harboring mutations in Casp1/11 [38]. Wild-type *S. Typhimurium* infection markedly reduced macrophage migration into the scratch, whereas a SPI-2-deficient mutant (*ΔssaV*) did not (Figs 1A and 1B). Consistent with previous findings in dendritic cells [10], an SspH2-deficient strain (*ΔsspH2*) was also attenuated for inhibition of host cell migration (Figs 1A and 1B), confirming that SspH2 contributes to the SPI-2-dependent suppression of macrophage migration. In agreement with recent findings that *sspH2* gene deletion did not cause a proliferation defect in RAW264.7 cells [46], wild-type RAW264.7 cells infected with wild-type or *ΔsspH2* strains of *S. Typhimurium* had statistically indistinguishable bacterial loads 24 hours post-infection (S1 Fig).

**Fig 1.**
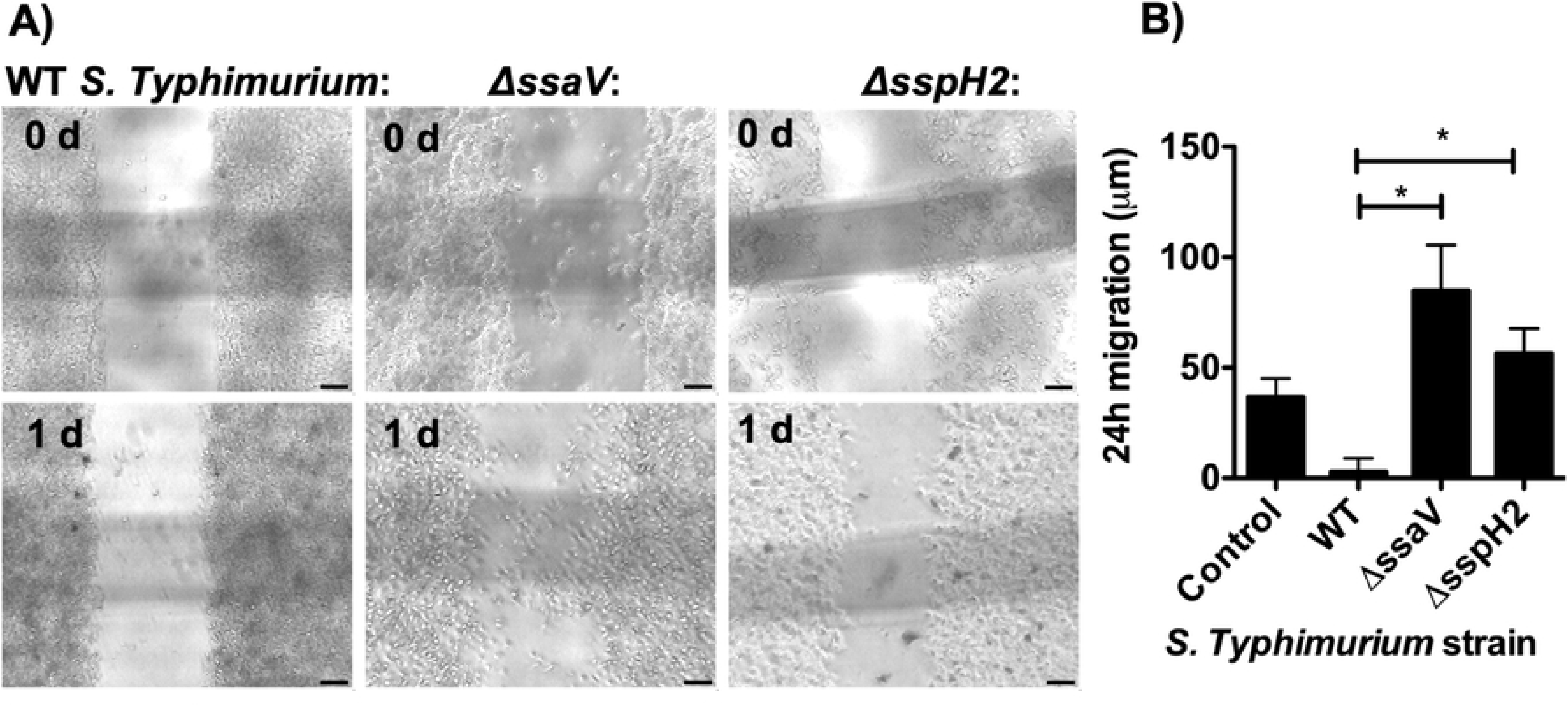
*S. Typhimurium* inhibits macrophage migration via SPI-2 and SspH2. Caspase 1/11 -/-RAW264.7 cells were infected with WT (SL1344) or the indicated mutant strain of *S. Typhimurium* at an MOI of 30:1 (mock-infected samples were used as a control). Immediately after infection, the monolayer was scratched with a pipette tip and micrographs were recorded under 100X magnification immediately and one day after infection. The width of every scratch at three specific points was calculated for each time point using Fiji software, and the migration distance was calculated as the difference between the scratch width at day zero and day one. Representative images are shown (A, scale bar = 100 µm). N ≥ 3, means and standard errors of the means (SEM) are presented in B, * p-value < 0.05 in a two-sample two-tailed Student’s t test.

### Conditional expression of SspH2 is sufficient to inhibit macrophage migration

To dissect SspH2’s specific contributions to this phenotype, we engineered a RAW264.7 cell line with doxycycline-inducible expression of SspH2-GFP. The *sspH2* coding sequence (with the stop codon replaced by *gfp*) was cloned downstream of a promoter containing a tetracycline response element and transduced into RAW264.7 cells stably expressing the reverse tetracycline transcriptional activator (rtTA), allowing tunable induction of SspH2-GFP upon doxycycline addition [39].

Conditional SspH2 expression significantly inhibited wound closure in the scratch assay (Fig 2A, representative images shown in Fig 2B). Time-lapse microscopy revealed that SspH2 expression reduced mean cell velocity (Fig 2C) and decreased the number of directional changes per cell track (Fig 2D), indicating that SspH2 inhibits cell motility. In contrast, conditional expression of SseI-GFP increased both cell velocity and turning frequency, consistent with prior reports that SseI promotes excessive reorientation rather than impeding motility [9]. Moreover, SspH2 expression did not impair chemotaxis toward the macrophage chemoattractant CCL5 (Fig 2F). Together, these findings demonstrate that SspH2 acts independently of chemotaxis pathways to inhibit the fundamental motility machinery of host cells.

**Fig 2.**
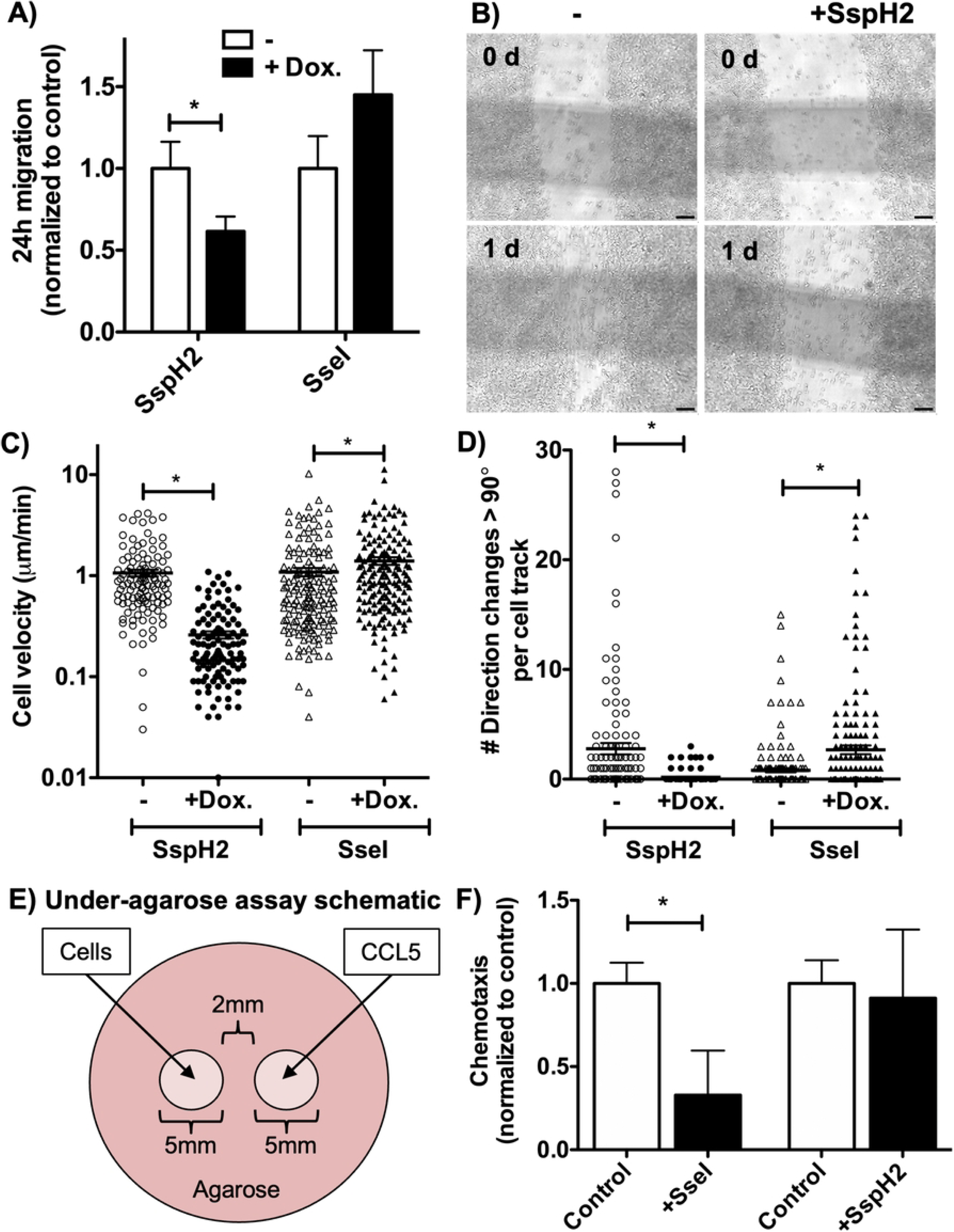
SspH2 expression alone is sufficient to impair macrophage migration. RAW264.7 cells were genetically modified to express the indicated SPI-2 effectors tagged with GFP under a doxycycline-inducible promoter. A-B) Scratches were introduced into the RAW264.7 monolayers with a 200µl pipette tip and treated with or without doxycycline (10 ng/mL for SspH2- and 100 ng/mL for SseI-expressing cells). Micrographs were recorded under 100X magnification immediately and 1 day later. Average migration (normalized to controls without doxycycline treatment) for at least 3 separate experiments is shown in A (error bars represent the standard deviation), and representative micrographs are shown in B (scale bar = 100 µm). C-D) SPI-2 effector-expressing RAW264.7 cells (as described in A and B) were seeded onto coverglass and treated with 0 or 10 ng/mL doxycycline for one day before observing by time-lapse microscopy every 3 minutes over 3.5 hours at 600X. Cells were tracked using Fiji to measure cell migration velocity (C) and the number of times a cell changes its direction of movement by more than 90 degrees (D). Between 113 and 163 cells were tracked for each group, and the mean and SEM are indicated by horizontal lines and error bars, respectively. E) A schematic of the under-agarose assay to measure chemotaxis: darker pink represents agarose while lighter pink represents wells (holes) cut out of the agarose where either cells or 500 ng/mL CCL5 was added. The indicated SPI-2 effector-expressing RAW264.7 cells were seeded into the well designated for cells in a 1:1 mixture with RAW264.7 cells genetically modified to express mCherry alone as an internal control. In the presence of 10 ng/mL doxycycline, chemotaxis was measured as the number of SPI-2 effector-expressing GFP+ cells that migrate under the agarose toward CCL5 minus the number of cells that migrate away. This raw measurement was then normalized to the raw chemotaxis measurements of the mCherry+ internal control. F) The indicated SPI-2 effector-expressing cell lines were treated with doxycycline and allowed to chemotax under the agarose for 24 hours before taking images at 600X on either side of the well containing cells. Chemotaxis measurements normalized to the internal control are presented. The means of between ≥ 5 replicates are shown. All * indicate p-values < 0.05 in two-sample two-tailed Student’s t tests.

### SspH2 localizes to F-actin-rich regions and reduces F-actin abundance

Previous studies showed that SspH2 localizes to the plasma membrane via S-palmitoylation [19] and interacts with cytoskeletal regulators such as α-filamin and profilin-1 [20, 21] and potentially myosin family proteins [22]. Using confocal microscopy, we confirmed that SspH2-GFP localizes predominantly to the plasma membrane of RAW264.7 cells and co-localizes with phalloidin-stained F-actin (Figs 3A and 3B) whereas mCherry alone does not (Fig 3C). Notably, total phalloidin signal intensity was reduced in cells expressing SspH2 (Fig 3D), suggesting that SspH2 alters the actin cytoskeleton.

**Fig 3.**
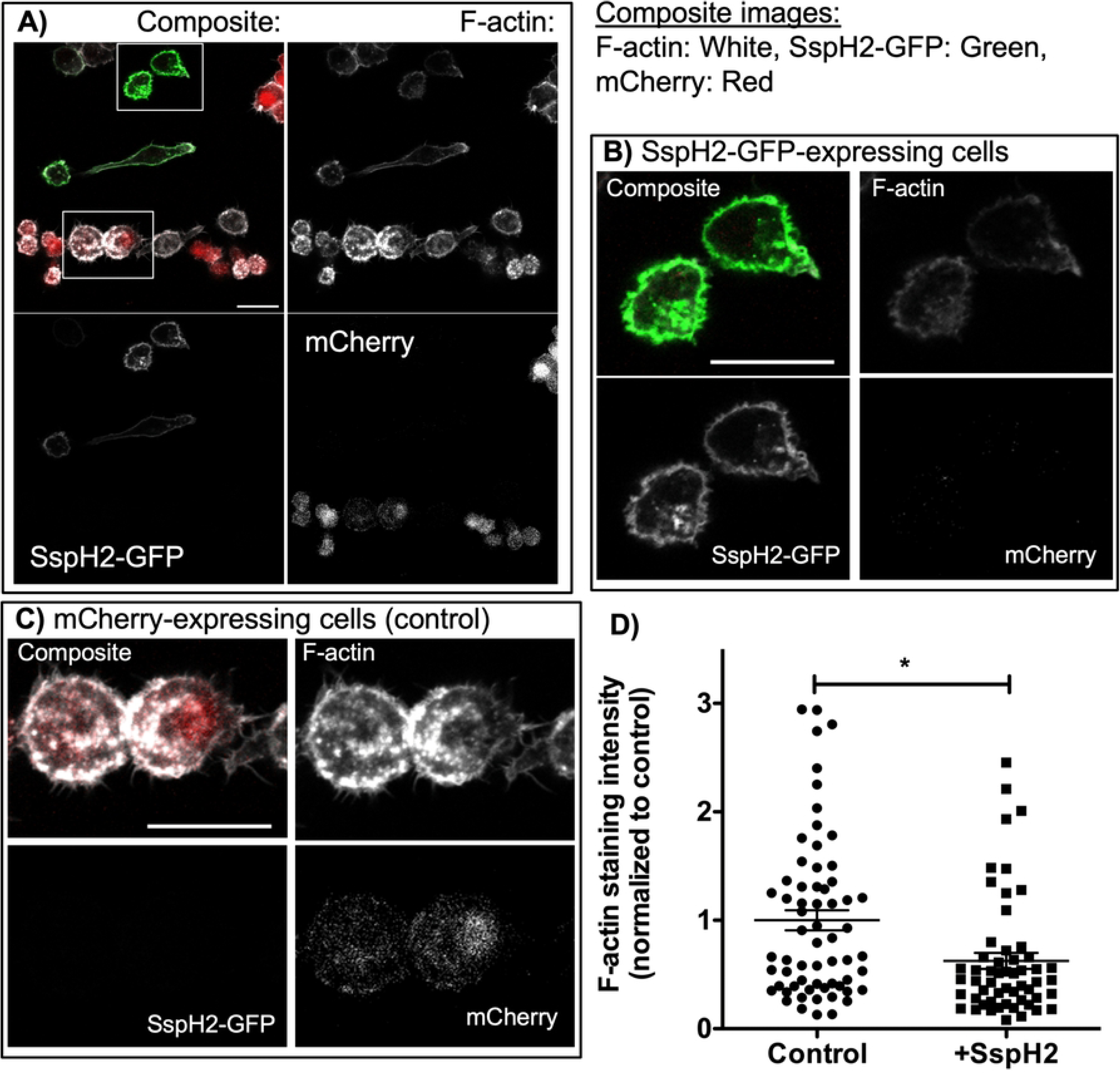
SspH2 localizes to the plasma membrane and reduces F-actin abundance in fixed cells. RAW264.7 cells genetically modified to express SspH2-GFP or mCherry alone (as described in Fig 2) were mixed together along with the parental cell line and seeded onto coverglass, then treated with doxycycline to induce SspH2-GFP expression. The next day, cells were fixed using paraformaldehyde and stained for F-actin using phalloidin-660. Confocal images were taken at 630X magnification and a representative image (z-slice) is shown in (A) with the boxed regions enlarged for two SspH2-GFP expressing cells in (B) and two control cells in (C) (scale bar = 21 µm). The total F-actin-associated fluorescent signal for each entire cell with and without SspH2-GFP expression was measured using Volocity software and presented in (D). At least 55 cells were assessed for each group, means ± SEM are indicated by horizontal line and error bars and * indicates p-value < 0.05 in a two-sample two-tailed Student’s t test.

To visualize actin dynamics in live cells, we co-expressed LifeAct-7-mCherry [47] in the SspH2-inducible cell line. Time-lapse video microscopy revealed a significant decrease in F-actin upon SspH2 induction (Figs 4A, 4Cii and 4Cv, S2 Video and S3 Video), confirming that SspH2 expression diminishes F-actin abundance in living macrophages. Without SspH2 expression, F-actin signal remained high and often condensed into bright punctate structures, sometimes arranged into rosette/ring-like formations (Fig 4Biv, arrow). Although these puncta were not further characterized by marker staining, their morphology is consistent with podosomes—F-actin-rich adhesive structures that play a key role in macrophage migration [29].

**Fig 4.**
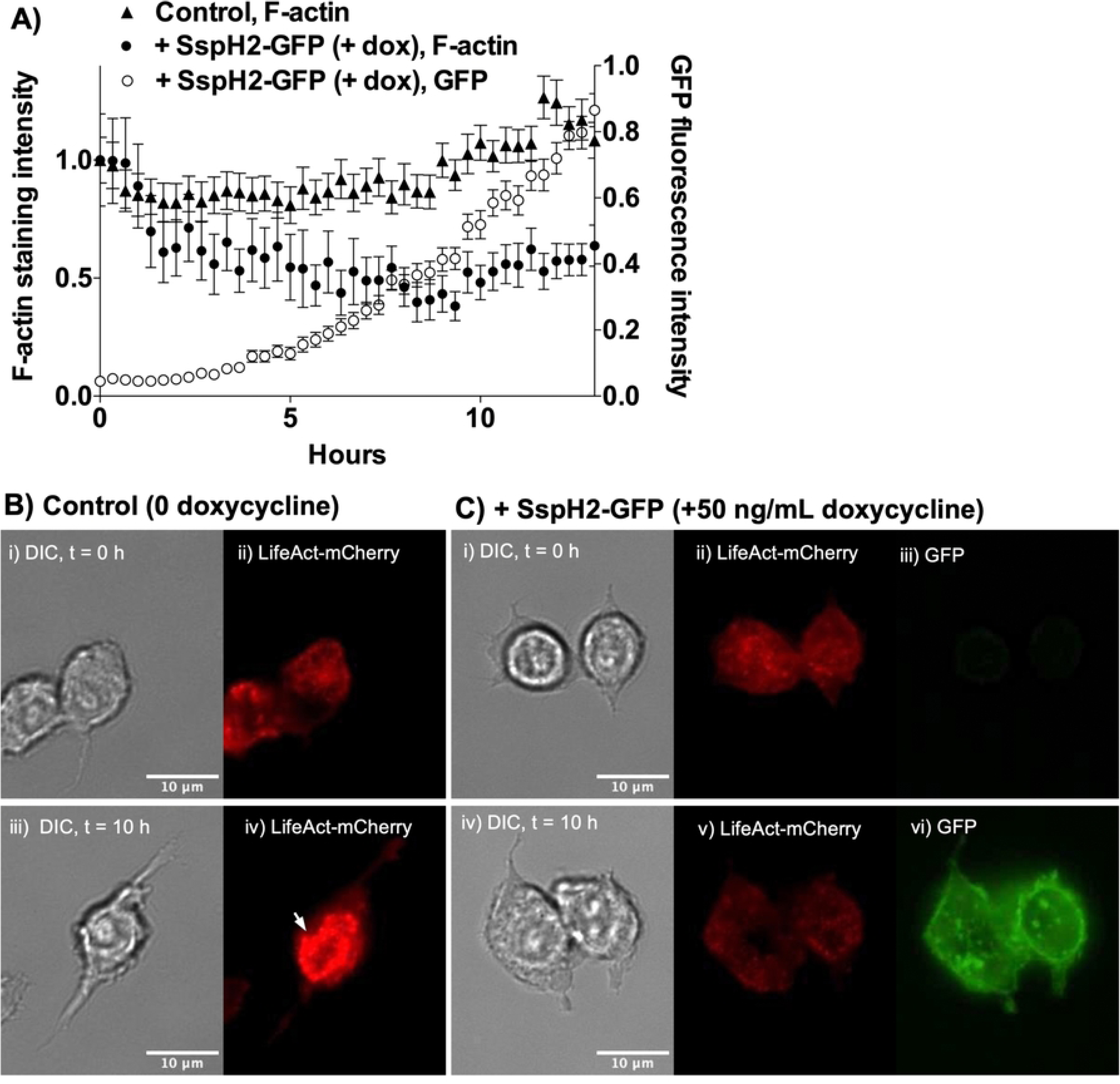
SspH2 disrupts F-actin in live cells. A) SspH2-GFP-expressing RAW264.7 cells were transduced with a constitutive expression construct for LifeAct-mCherry that stains F-actin in live cells. Cultures were treated with (circles) or without (control, triangles) doxycycline, and fluoresecent and DIC images were taken every 20 minutes over 13 hours at 600X magnification. CellTK software was used to track individual cells and to measure the fluorescence intensities of mCherry (filled symbols) and GFP (empty symbols) for each individual cell at each time point. The means ± SEM are shown, the number of cells for each time point ranged between 37 (at early time points) and 139 (at later time points). Control F-actin intensities were significantly different from that of cells treated with doxycycline for all times > 6 h (p < 0.05) in two-sample two-tailed Student’s t tests. Representative videos are shown in S2 Video (0 doxycycline) and S3 Video (+doxycycline). B-C) DIC and fluorescent images of representative cells are shown after addition of 0 (B, “Control”) or 50 ng/mL (C, “+SspH2-GFP”) doxycycline at 0 (Bi-ii, Ci-iii) or 10 h later (Biii-iv, Civ-vi). F-actin was detected using LifeAct-mCherry (red, Bii,iv and Cii,v) and SspH2 using GFP (green, Ciii,vi). Arrow points to an example of F-actin-rich punctate structures arranged into a rosette/ring formation.

### SspH2’s E3 ubiquitin ligase activity is required to disrupt the actin cytoskeleton

SspH2 contains a conserved catalytic cysteine residue (C580) required for E3 ubiquitin ligase activity [18]. To test whether enzymatic activity underlies cytoskeletal disruption, we generated an inducible RAW264.7 line expressing a catalytic mutant of SspH2 (SspH2C580A). In cells co-expressing LifeAct-7-mCherry, induction of SspH2C580A failed to reduce F-actin signal intensity (Fig 5A) as well as decrease cell migration velocity (Fig 5B), demonstrating that E3 ligase activity is essential for SspH2 to disrupt F-actin and to inhibit cell migration.

**Fig 5.**
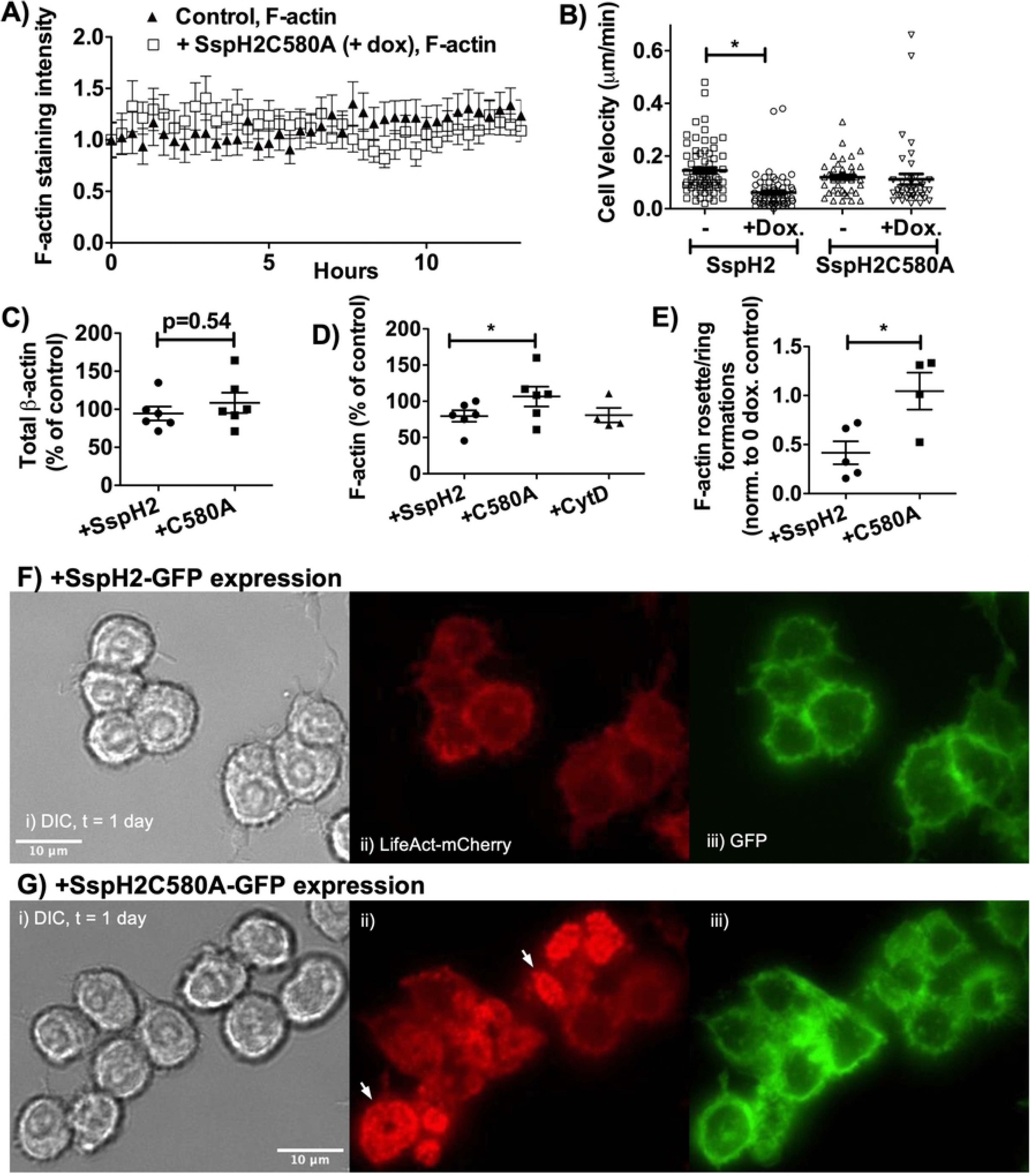
E3 ubiquitin ligase activity of SspH2 is required for cytoskeletal disruption. A) The *sspH2* gene was mutated to introduce an alanine in place of the catalytic cysteine residue at amino acid position 580 creating the catalytic mutant SspH2C580A. As described for Fig 4A, RAW264.7 cells were genetically modified to conditionally express SspH2C580A-GFP upon addition of doxycycline and to constitutively express the F-actin staining protein LifeAct-mCherry. Cultures were treated with (empty squares) or without (control, filled triangles) doxycycline, and then DIC and fluorescent images were taken every 20 minutes at 600X magnification. CellTK software was used to track individual cells and to measure the mCherry fluorescence intensities for each individual cell at each time point. The means ± SEM for each time point are shown, the number of cells for each time point was between 45 (at early time points) and 78 (at later time points). Excluding time = 8.67 h, control F-actin intensities were **not** significantly different from that of cells treated with doxycycline for all times shown (p > 0.05) in two-sample two-tailed Student’s t tests. B) After 12 h treatment with or without doxycycline, cells expressing SspH2 or the catalytic mutant SspH2C580A were tracked and the cell velocities were measured using Fiji. Between 41 and 69 cells were tracked for each group, the means ± SEM are shown, and * p-value < 0.05 in a two-sample two-tailed Student’s t test. C) The total β-actin was harvested from the indicated SspH2-expressing RAW264.7 cell lines (C580A = SspH2C580A mutant) treated with and without doxycycline (for one day) and quantified by western blot (normalized to GAPDH). The amount of total β-actin in doxycycline-treated samples was reported as a percent of samples without doxycycline treatment (control), and the p-value was calculated using a paired two-tailed Student’s t test. D) The fraction of β-actin existing as F-actin was quantified for the indicated SspH2-expressing RAW264.7 cell lines with and without doxycycline treatment for one day. As a positive control, RAW264.7 cells were treated for one day with 10 µM cytochalasin D (CytD), an inhibitor of actin polymerization that diminishes the cell’s F-actin fraction. The data for doxycycline-(or cytochalasin D-) treated samples are presented as the percent of non-treated samples (controls). * indicates p-value < 0.05 in a paired two-tailed Student’s t test. E) The frequency of cells containing rosette/ring-like formations of F-actin puncta after one day with doxycycline treatment was quantified for ≥ 4 replicates for each group, and then normalized to that of cells not treated with doxycycline. * p-value < 0.05 in a two-sample two-tailed Student’s t test. Images of SspH2-GFP-(F) or SspH2C580A-GFP-(G) expressing cells were taken one day after addition of doxycycline using DIC (i) and fluorescence (ii - iii), scale bar = 10 µm. To discern higher-order F-actin structures, images with similar and easily visible F-actin signal are shown here. Arrows indicate punctate F-actin-rich (podosome-like) structures arranged into rosette/ring-like formations.

In addition to microscopy-based methods, cell fractionation studies were carried out to measure the relative amounts of filamentous F-actin and monomeric G-actin in the cytosolic extracts of cells expressing either SspH2 or the catalytic mutant. Expression of SspH2 specifically impacted F-actin as total amounts of β-actin did not change (Fig 5C). Only catalytically active SspH2 significantly lowered the F-actin fraction, confirming that SspH2 E3 ligase activity is necessary for disruption of F-actin organization (Fig 5D).

### SspH2 disrupts higher-order F-actin structure

LifeAct-7-mCherry imaging revealed distinct punctate F-actin-rich structures in RAW264.7 cells, morphologically consistent with podosomes commonly detected in these cells [48, 49]. Previous work showed that disruption of podosomes correlated with decreased macrophage migration on 2-dimensional surfaces [50, 51], and reduction of higher-order structures like podosome rosettes correlated with reduced cell migration into dense 3-dimensional matrices [52], underlining the key role these F-actin-rich structures play in macrophage migration. In our study, these F-actin-rich puncta also could be found arranged in higher-order structures in the shape of rosettes/rings (Fig 4Biv, 5Gii). Expression of catalytically active SspH2 led to significantly fewer cells with these higher-order F-actin structures (Figs 5E and 5Fii), suggesting that SspH2 disrupts higher-order F-actin organization that may be essential for cell motility.

### SspH2-dependent inhibition of macrophage migration requires the presence of vinculin

Podosomes are composed of an F-actin-rich core surrounded by a complex of adhesion plaque proteins that anchor cells to the extracellular matrix via integrins [53]. Vinculin, a central component of these complexes, links actin to integrin-based adhesions as part of the “molecular clutch” that transduces traction forces at the leading edge [31, 54]. Cells lacking vinculin show impaired podosome rosette formation and migration in dense extracellular matrices [55–57].

CRISPR-targeting of the vinculin gene inhibited cell migration similarly to SspH2 expression (Fig 6A) and markedly reduced vinculin protein expression (Fig 6B). SspH2 expression did not further inhibit cell migration when vinculin expression was reduced (Fig 6A), indicating that removal of this host factor has rendered these cells insensitive to SspH2’s effect on cell migration. Re-expression of wild-type vinculin in the knockdown line increased baseline migration speed and restored the ability of SspH2 to inhibit migration (Fig 6C), confirming that the lack of SspH2’s inhibitory effect in the vinculin knockdown cells was due specifically to the reduced expression of vinculin. These findings demonstrate that the presence of normal levels of vinculin is necessary for SspH2-mediated suppression of host cell migration, suggesting that this effector targets vinculin-sensitive F-actin structures to interfere with macrophage motility.

**Fig 6.**
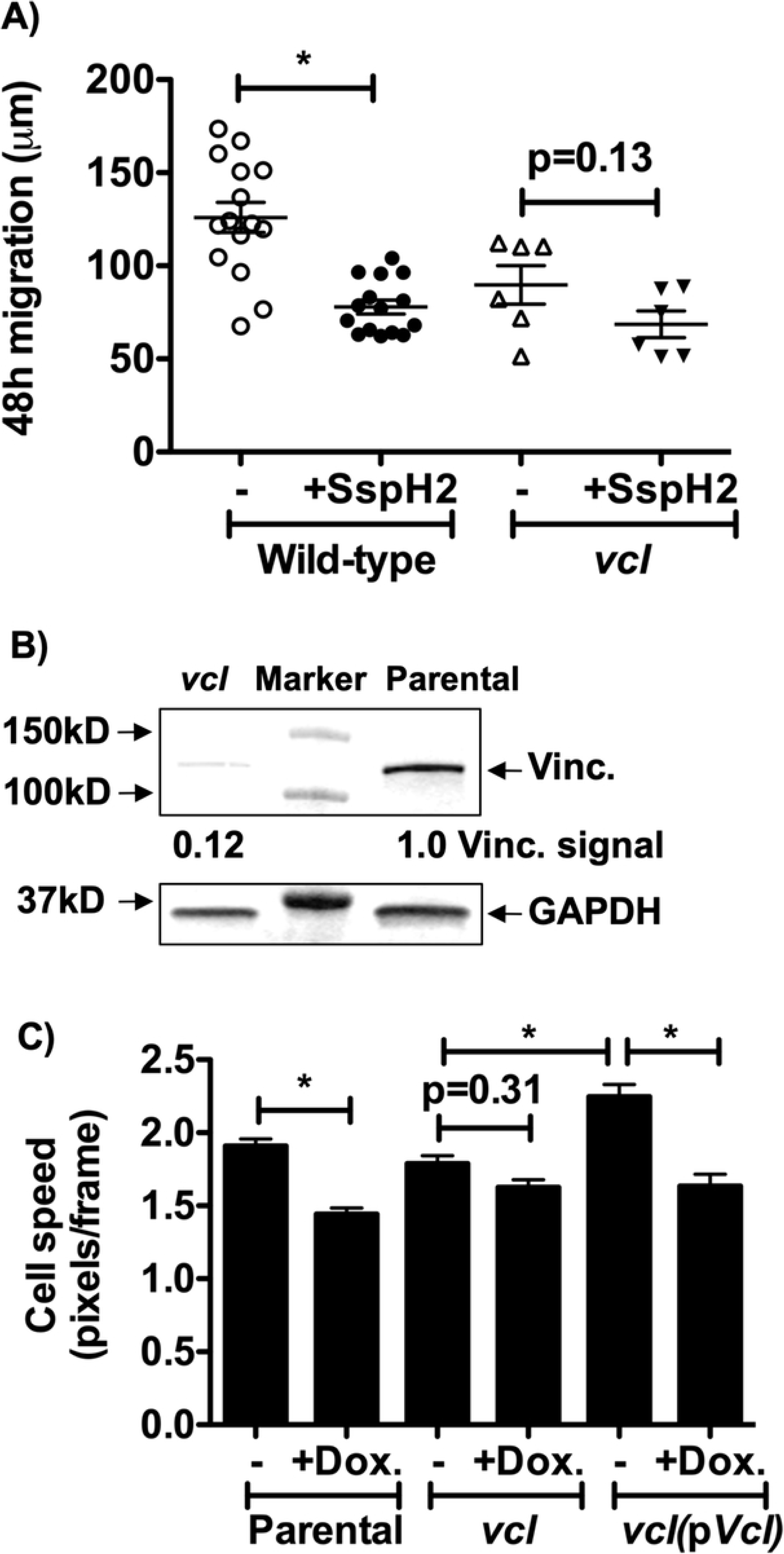
SspH2-dependent inhibition of cell migration requires the presence of vinculin. SspH2-GFP-expressing RAW264.7 cells (“parental” cell line) were transfected with an expression construct that co-expressed Cas9 and a small scaffold RNA targeting *vcl*, after which stable transfectants were selected. A) Scratches were introduced into confluent monolayers of the parental cell line (expressing wild-type vinculin) or cells stably expressing the small scaffold RNA targeting *vcl*, and treated with doxycycline to induce SspH2 (“-“ = 0 doxycylcline). The width of every scratch at three specific points was measured immediately after making the scratch and then again 48 hours later. Means ± SEM are shown for ≥ 6 replicates for each group, and * p-value < 0.05 in a two-sample two-tailed Student’s t test. B) Vinculin protein expression was assessed by western blot for the parental and *vcl*-targeted cell line. Vinculin signal was calculated as the vinculin intensity divided by the GAPDH intensity (and normalized to the value of the parental cells). C) A vinculin expression construct (p*Vcl*) was used to complement the *vcl* mutant phenotype, and the indicated cell lines were subjected to time-lapse microscopy, taking images every 20 minutes for 12 hours at 600X magnification. Cells in these time-lapse videos were then tracked using Fiji’s Trackmate plugin software (finding at least 695 cell tracks per group) and the straight line speed is presented (means ± SEM are shown, * p-value < 0.05 in a Tukey-Kramer multiple comparisons test, using JMP statistical software).

## Discussion

The *Salmonella enterica* SPI-2 effector SspH2 was first discovered to interfere with host cell migration in a genetic screen for mutants unable to block dendritic cell migration [10]. Here, we confirm that *S. Typhimurium ΔsspH2* is attenuated in its ability to inhibit macrophage migration in vitro (Fig 1) and show that SspH2 expression alone is sufficient to recapitulate this phenotype in RAW264.7 cells (Fig 2). In addition to reducing migration speed, SspH2 expression decreased F-actin (Figs 3 and 4) and the number of cells with higher-order F-actin-rich structures arranged into rosette/ring-like formations (Fig 5), implicating cytoskeletal remodeling as the underlying mechanism. SspH2-dependent inhibition of host cell migration was sensitive to the presence of the host podosome protein, vinculin (Fig 6), suggesting a potential relationship between SspH2 and these F-actin-rich host structures. Vinculin plays an essential role in focal adhesions and podosomes, functioning as part of a “molecular clutch” that couples F-actin polymerization to membrane protrusion at the leading edge of migrating cells [31–33, 54]. Accordingly, deletion of the *vcl* gene impairs podosome rosette/ring formation in osteoclasts [57] and alters cell motility and adhesion dynamics in murine embryonic fibroblasts [55, 56, 58]. Considering that effectors from other intracellular pathogens directly target vinculin, this factor may represent a convenient host node that pathogens use to disrupt normal cell movement behavior, favoring colonization and persistence.

SspH2 belongs to the NEL family of bacterial E3 ubiquitin ligases, whose members are found across diverse genera, including *Shigella*, *Salmonella*, *Edwardsiella*, *Bradyrhizobium*, *Rhizobium* and *Pseudomonas* [59]. NEL family effectors such as *Shigella* IpaH [60, 61] and *Yersinia* YopM [62] are required for virulence in animal models, and catalytic mutants fail to complement the mutant phenotypes. Similarly, the catalytically inactive SspH2C580A mutant did not alter F-actin organization or cell motility in RAW264.7 cells, confirming that SspH2’s E3 ubiquitin ligase activity is essential for its function (Fig 5). Knockdown of vinculin expression mimicked SspH2’s inhibitory effect on cell migration, and without normal levels of vinculin, SspH2 expression did not further inhibit cell migration (Fig 6). While these data indicate that SspH2 may act in a pathway that converges on an F-actin regulatory mechanism involving vinculin, it remains to be determined whether vinculin is a direct target of SspH2 or acts downstream on another substrate. Although vinculin itself has not been reported as a target of mammalian E3 ubiquitin ligases, other vinculin-associated adhesion plaque proteins such as talin [63] and paxillin [64, 65] are ubiquinated, and dysregulation of these factors leads to altered cell migration. Future work will be necessary to determine whether SspH2 modifies vinculin directly or perturbs a vinculin-associated pathway, possibly involving LMO4 or other host targets linked to cytoskeletal control.

Manipulation of host cell migration is an emerging and unifying strategy among intracellular bacterial pathogens. We previously demonstrated that the SPI-2 effector SseI (SrfH) inhibits macrophage and dendritic cell migration and promotes persistent *S. Typhimurium* infection in mice [9]. This phenotype requires SseI’s deamidase activity [13] and IQGAP1 [9], a host scaffolding protein that regulates cytoskeletal dynamics. Wild-type *S. Typhimurium* exhibits a competitive advantage over *ΔsseI* mutants in vivo, underscoring the importance of migration inhibition for persistence in the host [9]. Similarly, *Legionella pneumophila* suppresses macrophage and epithelial cell migration via its Icm/Dot type IV secretion system and the quorum sensing molecule LAI-1, which acts through IQGAP1 [43, 66]. *Chlamydophila (Chlamydia) pneumoniae* also regulates vascular smooth muscle cell migration via an IQGAP1-dependent mechanism [67]. The convergence of these diverse intracellular pathogens on IQGAP1 and other cytoskeletal regulators highlights that control of host cell migration is a conserved virulence strategy.

Beyond IQGAP1, multiple bacterial pathogens deploy effectors that directly modulate adhesion and cytoskeletal components. The *Chlamydia trachomatis* effector TarP binds vinculin, remodeling focal adhesions and inhibiting cell motility [37, 42, 68]. *Shigella flexneri* IpaA targets vinculin leading not only to invasion but also to increased cell adhesion [34, 35]. The *Shigella* effector OspE targets integrin-linked kinase (ILK) also to increase cell adhesion [36], while the effector IpgD disrupts chemotaxis by depleting PIP_2_ levels in immune cells [69]. *Yersinia enterocolitica* YopT, a cysteine protease, cleaves Rho GTPases and disrupts F-actin-rich podosomes key for robust macrophage migration [70]. Taken together, these findings highlight a recurring mechanistic theme whereby bacterial pathogens use translocated effectors to target host factors, such as vinculin, IQGAP1 or G proteins, that control F-actin dynamics and cell migration.

Our findings identify SspH2 as a new player in this broader strategy. By co-opting ubiquitin-dependent regulation of the actin cytoskeleton, SspH2 limits macrophage migration likely by targeting vinculin-sensitive F-actin-rich structures. This work expands the known repertoire of *Salmonella* virulence strategies that modulate host cell behavior and raises intriguing questions about how migration inhibition contributes to immune evasion and long-term persistence in vivo. Future studies will be needed to define the exact host substrates SspH2 targets to manipulate host cell migration and to determine whether this effector’s impact on F-actin organization promotes bacterial survival within infected tissues and ultimately persistent infection.

## Acknowledgments

We acknowledge Denise Monack for providing expertise with respect to murine cell models of *Salmonella* infection, access to time-lapse microscopy equipment, laboratory space and reagents necessary for completion of this project. We also acknowledge Manuel Amieva for providing expertise with respect to F-actin cytoskeleton dynamics and confocal microscopy, laboratory space and reagents necessary for this project. We also acknowledge Trung Hoang Minh Pham for lending his flow cytometry expertise and Cole Dovey for sharing the pLenti-mCherry vector with us. Finally, we would like to acknowledge the comprehensive support provided by the administrative staff, professors and students of Holy Names University, without whom none of this research would have been possible.

## Supporting information

**S1 Fig. Bacterial loads of WT and *ΔsspH2 S. Typhimurium* in RAW264.7 cells are equivalent.** RAW264.7 cells were infected with GFP-expressing wild-type (WT) or *ΔsspH2 S. Typhimurium* strains and DIC and fluorescent images were taken 24 hours later at 600X magnification. The fraction of cells containing fluorescent bacteria was quantified using Fiji and the means ± SEM are shown (N = 3).

